# EOGT Enables Residual Notch Signaling in Mouse Intestinal Cells Lacking POFUT1

**DOI:** 10.1101/2023.04.14.536788

**Authors:** Mohd Nauman, Shweta Varshney, Jiahn Choi, Leonard H. Augenlicht, Pamela Stanley

**Author notes:** Correspondence: Pamela Stanley, Dept. Cell Biology, Albert Einstein College of Medicine, 1300 Morris Park Ave., New York, NY 10641.

## Abstract

Notch signaling determines cell fates in mouse intestine. Notch receptors contain multiple epidermal growth factor-like (EGF) repeats modified by O-glycans that regulate Notch signaling. Conditional deletion of protein O-fucosyltransferase 1 (*Pofut1*) substantially reduces Notch signaling and markedly perturbs lineage development in mouse intestine. However, mice with inactivated *Pofut1* are viable, whereas complete elimination of Notch signaling in intestine is lethal. Here we investigate whether residual Notch signaling enabled by EOGT permits mice lacking *Pofut1* in intestine to survive. Mice globally lacking *Eogt* alone were grossly unaffected in intestinal development. In contrast, mice lacking both *Eogt* and *Pofut1* died at ∼28 days after birth with greater loss of body weight, a greater increase in the numbers of goblet and Paneth cells, and greater downregulation of Notch target genes, compared to *Pofut1* deletion alone. These data establish that both O-fucose and O-GlcNAc glycans are fundamental to Notch signaling in the intestine and provide new insights into roles for O-glycans in regulating Notch ligand binding. Finally, EOGT and O-GlcNAc glycans provide residual Notch signaling and support viability in mice lacking *Pofut1* in the intestine.

## Introduction

The small intestine of male and female mammals is comprised of differentiated epithelial lineages derived from *Lgr5^hi^*-expressing and potentially other intestinal stem cells (ISC), which maintain overall mucosal homeostasis and provide principal functions in nutrient uptake and metabolism ^1, 2^. Notch signaling controls the functioning of ISCs by regulating maintenance of the stem cell pool, and specifying lineage differentiation to absorptive enterocyte cells or secretory goblet, Paneth, enteroendocrine and Tuft cells ^3^. The critical role of Notch signaling is mechanistically complex, however, dependent on post-translational modifications of Notch receptors and ligands that fine tune the strength of Notch signals known to be central in embryonic and tissue development ^4^. Fundamental to this fine tuning are epidermal growth factor-like (EGF) repeats present in the extracellular domain of Notch receptors (NECD) that are modified by O-glycans including O-glucose, O-fucose and O-GlcNAc glycans (Fig. 1*A*). These O-glycans regulate Notch signaling in cell-based assays, in mouse models and in humans ^4–7^. Only ∼100 proteins contain one or more EGF repeats with appropriate consensus sites for O-glycan addition ^8^, and not all O-glycans necessarily affect biological function ^9^. However, O-fucose on EGF12 of NOTCH1 interacts directly with Notch ligands DLL4 ^10^ and JAG1 ^11^ to facilitate Notch-ligand interactions and the loss of this fucose is embryonic lethal in the C56BL/6 background ^12^. Importantly, amongst proteins with modifiable EGF repeats, Notch receptors contain by far the greatest number, and thus phenotypes arising from the loss of O-glycan glycosyltransferases commonly reflect defective Notch signaling ^4–7^.

**Figure 1.**
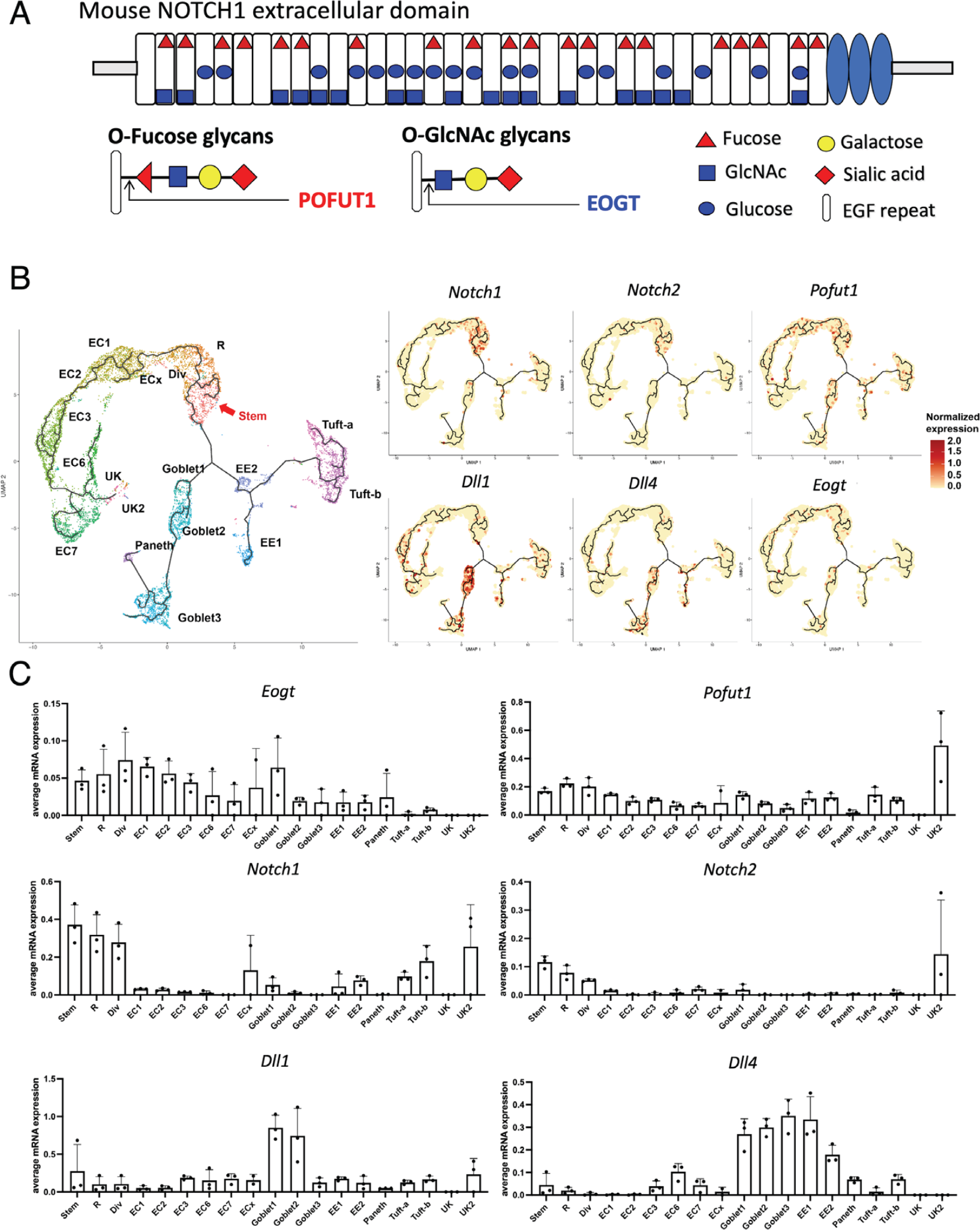
O-glycans on NOTCH1 and expression of *Eogt* and *Pofut1* in mouse intestine. (*A*) Diagram of mouse NOTCH1 ECD showing O-fucose and O-GlcNAc glycans predicted at EGF repeats that contain the appropriate amino acid consensus sequence. Each O-glycan (including O-glucose glycans) is indicated by the initiating sugar. (*B*) *Eogt* and *Pofut1* expression in mouse intestinal cells by scRNAseq. The trajectory analysis of the combined data from intestinal cells of 3 mice is shown. The definition of each intestinal cell type is given in Abbreviations. (C) Transcript levels of candidate genes expressed in each cell type were quantitated for 3 mice.

In the intestine, conditional deletion (cKO) by Villin-Cre of *Pofut1,* the gene encoding the enzyme that adds O-fucose to appropriate EGF repeats, causes a substantial reduction in Notch signaling and a marked increase in secretory cell lineages in male and female mice ^13^. *Pofut1* cKO mice are smaller than littermates but survive for at least 6 months. Mice lacking a single Fringe gene (*Lfng* or *Rfng*) encoding enzymes that add GlcNAc to the O-fucose transferred to EGF repeats by POFUT1, also exhibit altered Notch signaling and a Notch-defective intestinal phenotype ^14^. A clear illustration of the importance of modulating Notch signaling, rather than Notch acting as an “on-off switch”, is that mice with down-regulation of Notch by cKO of *Pofut1* survive for ≥ 6 months on a mixed genetic background ^13^, whereas cKO of two Notch ligand genes *Dll1* and *Dll4* targeted by Villin-Cre causes death at 4-6 days after the final tamoxifen dose ^15^. Similarly, conditional deletion of *Notch1* and *Notch2* ^16^, or treatment with a gamma-secretase inhibitor ^17^, produce severe inhibition of Notch signaling in the intestine.

Given the viability of *Pofut1*[F/F]:Villin-Cre mice, we hypothesized that residual Notch signaling may be enabled by O-GlcNAc glycans on Notch receptors. EOGT is the EGF domain specific GlcNAc transferase that initiates modification of EGF repeats by O-GlcNAc glycans ^18^. *Eogt* null pups exhibit altered development of the postnatal retina typical of defective Notch signaling, and EOGT regulates Notch receptor-ligand interactions ^19^. Therefore, a dual approach was used to investigate interactive roles of O-fucose and O-GlcNAc glycans on Notch receptors. First, Chinese hamster ovary (CHO) cells lacking *Eogt*, *Pofut1* or both *Eogt* and *Pofut1* were generated by a CRISPR/Cas9 strategy and examined for binding of canonical Notch ligands. Second, small intestine of male and female mice in which *Eogt* and *Pofut1* were genetically inactivated individually or in combination were interrogated. The data establish that, while mice lacking *Eogt* from conception did not exhibit an obvious intestinal phenotype, marked Notch signaling defects in *Pofut1* cKO intestine were significantly enhanced when both *Eogt* and *Pofut1* were inactivated. The combined data clearly demonstrate synergy between O-fucose and O-GlcNAc glycans in regulating Notch signaling in the intestine.

## Results

### *Eogt*, *Pofut1*, Notch Receptors and Ligands expressed in Mouse Intestine

Fine tuning of Notch signaling is fundamental in normal development and is achieved by post-translational modification of Notch receptors by glycosylation ^5–7^. Figure 1A illustrates consensus sites of predicted glycosylation of mouse NOTCH1 by O-fucose initiated by POFUT1, and by O-GlcNAc initiated by EOGT ^20, 21^, and the subsequent extension by different enzymes. O-fucose and O-GlcNAc glycans are recognized by Notch ligands whereas O-glucose glycans are not ^22^. Roles for Notch signaling have been clearly documented in intestinal stem cell functions ^15, 16^, but fine tuning in the intestine has not been defined. scRNAseq analysis for cells of the mouse intestinal epithelium identified complex relationships between expression of *Notch1* and *Notch2* receptor genes, the *Delta1* and *Delta4* Notch ligand genes and the *Eogt* and *Pofut1* glycosyltransferase genes (Fig. 1B and 1C). While *Notch1* and *Notch2* were expressed mainly in stem and progenitor cells, the genes encoding *Dll1* and *Dll4* were predominantly expressed in early goblet cells, with *Delta4* also expressed in early enterocytes. Importantly, *Pofut1* and *Eogt* were expressed in the same cell types as Notch receptors and ligands, also more heterogeneously. We therefore investigated further roles for Notch signaling in intestinal homeostasis by genetic inactivation of *Eogt* and/or *Pofut1* to determine differential impacts on the differentiation of cells along the crypt-villus axis. But first we investigated effects of O-glycan removal on Notch ligand binding in a cell-based assay using CHO cells.

### Notch Ligand Binding to CHO O-glycan Mutants

To investigate potential synergism between O-fucose and O-GlcNAc glycans (Fig. 1A), deletion mutants were generated by a CRISPR/Cas9 strategy in Lec1 CHO cells ^23^. Lec1 CHO cells do not make hybrid or complex N-glycans and therefore provide a simplified glycosylation environment for glycosyltransferase gene deletions affecting O-glycan synthesis ^24^. Lec1 cells harboring inactivated *Eogt*, *Pofut1* or *Eogt* and *Pofut1* genes were characterized as described in Methods and Supplementary Figure S1. To determine if the cell surface expression of NOTCH1 (as an example of Notch receptors) was affected by the loss of O-glycan subsets from its extracellular domain (NECD1), Lec1 and CRISPR/Cas9 Lec1 mutants were assessed for binding of an anti-NECD1 antibody by flow cytometry. Modestly reduced expression of cell surface NOTCH1 was observed in *Eogt:Pofut1* dKO Lec1 cells (Fig. 2A and 2B). By contrast, the binding of soluble Notch ligands DLL1-Fc and DLL4-Fc was markedly reduced in both *Pofut1* and *Eogt:Pofut1* mutant Lec1 cells (Fig. 2A and 2B). The reductions in binding observed in *Eogt* KO compared to control cells and in *Eogt:Pofut1* dKO compared to *Pofut1* cKO cells were not statistically significant, but suggested potential synergism between O-fucose and O-GlcNAc glycans. This hypothesis was pursued *in vivo*.

**Figure 2.**
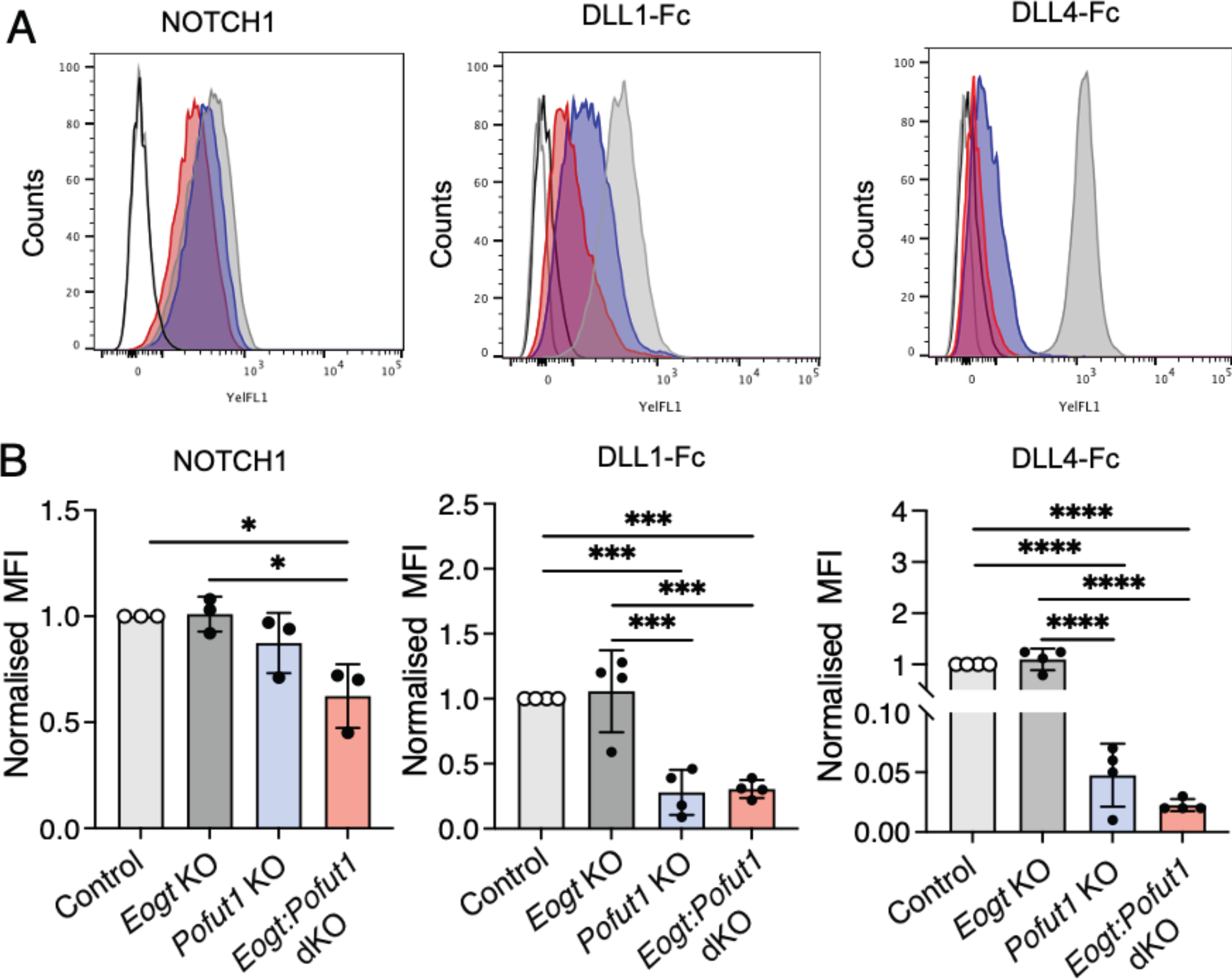
Notch ligand binding to Lec1 cells lacking EOGT, POFUT1 or both. *(A)* Representative flow cytometry profiles for anti-NOTCH1 ECD and Notch ligands DLL1-Fc and DLL4-Fc binding to Control (gray), *Eogt* KO, *Pofut1* KO (blue) and *Eogt:Pofut1* dKO (red) Lec1 cells. Secondary antibody alone for *Pofut1* KO (black line) and *Eogt:Pofut1* dKO (gray line). *(B)* Mean fluorescence index (MFI) for binding of anti-NOTCH1 ECD or different Notch ligands to *Eogt* KO, *Pofut1* KO or *Eogt:Pofut1* dKO CHO cells normalized to MFI of Control cells (n = 3-4 experiments). *P* values from one-way ANOVA followed by Tukey’s multiple comparisons test - \**P* < 0.05, \*\*\**P* < 0.001, \*\*\*\**P* < 0.0001.

### *Eogt* Supports Residual Notch Signaling in *Pofut1* cKO Intestine

Intestinal development was investigated in mice homozygous for *Eogt* inactivation compared to heterozygotes and wild type littermates. There were no differences in body weight, length of small intestine, production of secretory cells, or morphology and dimensions of villi or crypts in *Eogt*[+/+], *Eogt*[+/-] and *Eogt*[-/-] mice (Supplementary Figure S3). To investigate potential alterations at the molecular level, intestinal crypts were prepared by fractionation as described in Methods. The relative enrichment of *Lyz1* in crypts (fraction IV) and *Fabp2* in villi (fraction I) was determined by qRT-PCR (Supplementary Figure S4A and S4B). *Notch1* and *Lgr5* transcripts were reduced by homozygous inactivation of *Eogt* (Supplementary Figure S4C-S4G), indicating an impact of the loss of *Eogt* on *Lgr5*+ stem cells, shown to express *Notch1* in Figure 1B. However, these effects did not lead to altered expression of Notch signaling target genes such as *Hes1* in *Eogt* null crypts. Several attempts to perform scRNAseq on *Eogt*[-/-] EpCAM+CD45-cells sorted from crypt cells were not successful due to low cell viability. Finally, in all morphological, histological and molecular parameters examined, wild type and *Eogt*[+/-] intestine were indistinguishable, and thus *Eogt* heterozygous mice were appropriate controls for evaluating effects in mice with compound mutations in both *Eogt* and *Pofut1*.

Conditional inactivation of *Pofut1* in the mouse intestine greatly inhibits canonical Notch signaling, but the mice are viable and live to ≥ 6 months of age, albeit with significantly altered lineage representation ^13^. To test the hypothesis that EOGT and O-GlcNAc glycans provide residual Notch signaling in *Pofut1* cKO mice, we compared intestinal development in *Pofut1* cKO versus *Eogt:Pofut1* dKO mice. *Eogt*[-/-]*Pofut1*[F/F] and *Eogt*[+/-]*Pofut1*[F/+]:Villin-Cre mice were crossed to produce compound genotypes (Table 1). Whereas control and *Eogt*[-/-] pups from the first three litters were viable and developed without apparent defects, all *Eogt*:*Pofut1* dKO pups died at ∼P28. This was a strong indication that *Eogt* contributed to development in the *Pofut1* cKO intestine. In subsequent litters, histopathology of small intestine was assessed at P15. There were no changes in body weight, villi length or crypt depth in small intestine of *Eogt* KO, *Pofut1* cKO or *Eogt:Pofut1* dKO pups compared to control mice at P15 (Supplementary Figure S5A-S5C). The numbers of Alcian Blue-positive goblet cells and eosin-stained Paneth cells in the small intestine of *Eogt* KO mice were similar compared to controls (Supplementary Figure S5D-S5G). By contrast, goblet cells significantly increased in crypts and villi of *Pofut1* cKO and *Eogt:Pofut1* dKO pups (Supplementary Figure S5D-S5F). Importantly, the number of goblet cells was higher in *Eogt:Pofut1* dKO compared to *Pofut1* cKO intestine, indicating a contribution of *Eogt* to maintaining secretory cell development. Consistent with this, there was also an increase in Paneth cells in *Eogt:Pofut1* dKO crypts compared to *Pofut1* cKO crypts (Supplementary Figure S5G). Thus, while *Eogt* deletion had no apparent effects at P15, cKO of *Pofut1* caused a shift to greater secretory cell differentiation which was exacerbated by the concomitant elimination of *Eogt* in *Eogt:Pofut1* dKO pups.

**Table 1:** Eogt:Pofut1 dKO pups die at ∼28 dpp

| Mouse Group | Genotype | *Eogt* | *Pofut1* | Total Pups | Viability at ~P28 (%) |
| --- | --- | --- | --- | --- | --- |
| Control | *Eogt*<sup>+/-</sup>*Pofut1*<sup>F/+</sup><br>*Eogt*<sup>+/-</sup>*Pofut1*<sup>F/F</sup> | + | + | 14 | 100 |
| *Eogt* KO | *Eogt*<sup>-/-</sup>*Pofut1*<sup>F/+</sup><br>*Eogt*<sup>-/-</sup>*Pofut1*<sup>F/F</sup> | - | + | 9 | 100 |
| *Pofut1* cKO | *Eogt*<sup>+/-</sup>*Pofut1*<sup>F/F</sup>: Villin-Cre | + | - | 3 | 100 |
| *Eogt:Pofut1* dKO | *Eogt*<sup>-/-</sup>*Pofut1*<sup>F/F</sup>: Villin-Cre | - | - | 5 | 0 |

The cross *Eogt*<sup>-/-</sup>*Pofut1*<sup>F/F</sup> X *Eogt*<sup>+/-</sup>*Pofut1*<sup>F/+</sup>:Villin-Cre produced a range of different genotypes. Viability of progeny from the first 3 litters is shown.

Since *Eogt:Pofut1* dKO mice died about 1 month after birth, intestinal development was investigated at ∼P28. Body weight and length of small intestine were significantly decreased in *Eogt*:*Pofut1* dKO compared to *Pofut1* cKO mice (Fig. 3A and 3B). Both goblet and Paneth cells were significantly increased in *Eogt*:*Pofut1* dKO compared to *Pofut1* cKO intestine (Fig. 3C-3F). Villus length was decreased in *Eogt:Pofut1* dKO intestine compared to control, *Eogt* KO and *Pofut1* cKO intestine (Supplementary Figure S6A). Crypt depth, crypt length and crypt width were increased in *Pofut1* cKO and *Eogt:Pofut1* dKO compared to control or *Eogt* KO intestine (Supplementary Figure S6B and S6D). Thus, for all parameters assessed, conditional deletion of *Pofut1* from mice lacking *Eogt* in intestine caused a more severe phenotype than deletion of *Pofut1* alone,

**Figure 3.**
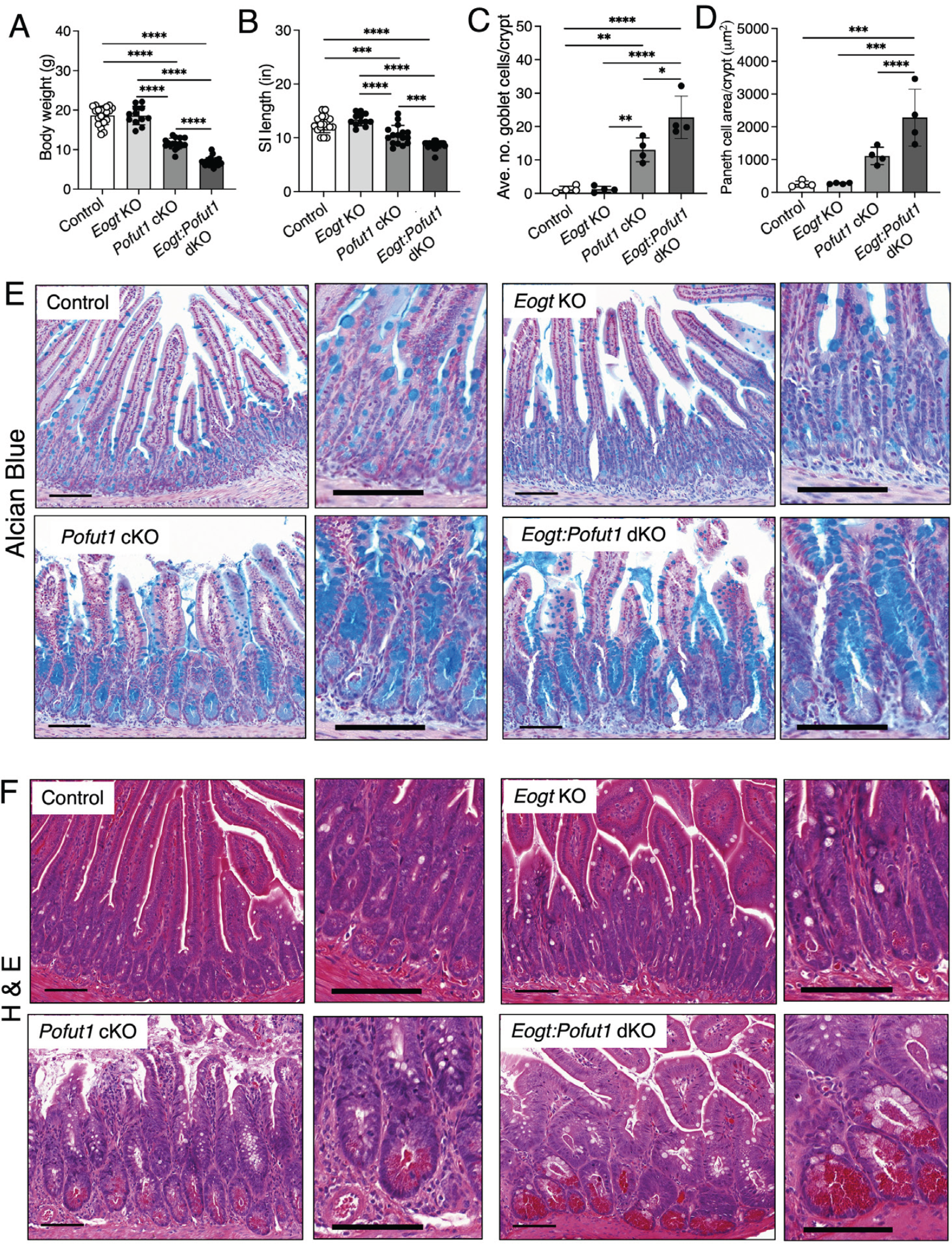
Synergy between *Pofut1* and *Eogt* in intestinal homeostasis. (*A* and *B*) Body weight and length of small intestine (SI) of Control, *Eogt* KO, *Pofut1* cKO and *Eogt:Pofut1* dKO mice (n ≥ 12 mice per group). (*C* and *D*) Number of goblet cells and area of Paneth cells in the crypts of experimental mice (∼50 crypts were analyzed in 4 mice per group). (*E* and *F*) Representative images showing goblet and Paneth cells in the small intestine of experimental mice (n = 4 mice per group). P values from one-way ANOVA followed by Tukey’s multiple comparisons test - *P < 0.05, **P < 0.01, ***P < 0.001, ****P < 0.0001.

### Notch Activation is Markedly Reduced in *Pofut1* cKO and *Eogt:Pofut1* dKO Crypts

Notch signaling is active at the base of crypts within Notch receptor-expressing ISC supported by ligand-expressing Paneth cells ^14^. Cleaved NICD1 (activated NOTCH1) is mainly detected in ISC and transit amplifying (TA) cells located in the crypt base ^25^, and our scRNAseq trajectory analysis of Notch receptor and ligand expression are consistent with this (Fig. 1B and 1C). To investigate Notch signaling status in crypt cells, NOTCH1 cleavage to NICD1 was determined by immunohistochemistry (Fig. 4A and 4B). Crypts of *Pofut1* cKO and *Eogt*:*Pofut1* dKO intestine were enlarged due to a significant increase in secretory cells and a concomitant reduction in crypt base columnar ISC. Val1744 antibody detects cleaved NOTCH1 and positive cells were rare to non-existent in both *Pofut1* cKO and *Eogt*:*Pofut1* dKO crypts (Fig. 4A). Control sections showed abundant signal for cleaved NOTCH1 in nuclei. Comparisons of signal for expression of cell surface NOTCH1 detected by antibody to NECD1 revealed no significant differences between *Pofut1* cKO and *Eogt*:*Pofut1* dKO, which were also similar to control (Fig. 4B). Specificity of signals for antibody to NICD1 and NECD1 was confirmed in sections stained with secondary antibody alone (Supplementary Figure S7A and S7B). Further, western blot analyses showed activated NOTCH2 in control crypt cells, but no signal for NICD2 cleavage in either *Pofut1* cKO or *Eogt*:*Pofut1* dKO crypt lysates (Fig. 4B).

**Figure 4.**
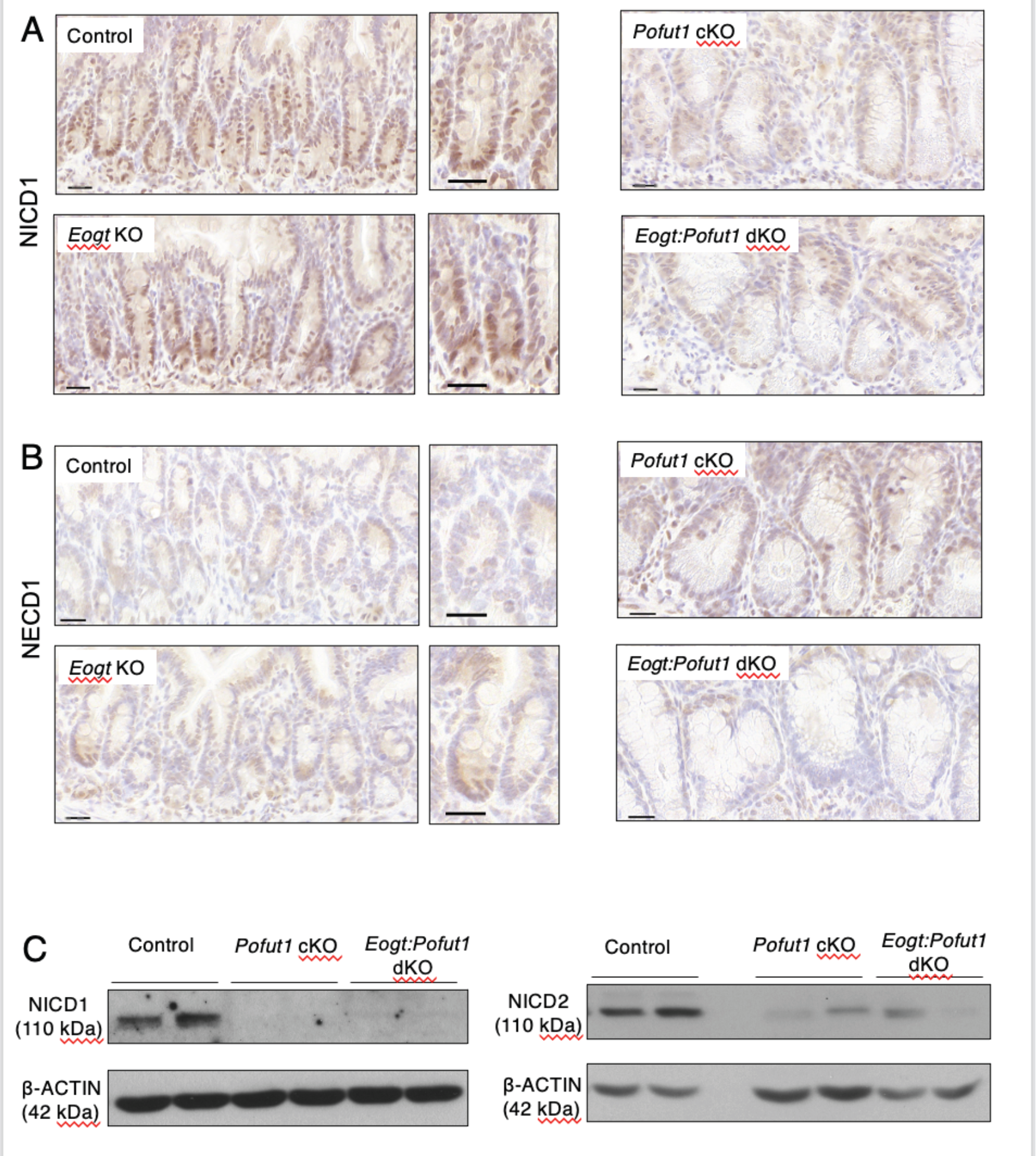
Expression of activated, cleaved NOTCH1 and NOTCH2 in intestine. (*A* and *B*) Immunohistochemistry of intestinal sections for binding of antibody to cleaved NOTCH1 (NICD1) or full length NOTCH1 (NECD1) in crypts of Control, *Eogt* KO, *Pofut1* cKO and *Eogt:Pofut1* dKO mice (n = 1-3 mice per group). (*C*) Western blots for cleaved NICD1 and NICD2 in the crypts of Control, *Pofut1* cKO and *Eogt:Pofut1* dKO (representative images of 4-6 mice per group).

### Notch Ligand Binding to *Eogt* KO, *Pofut1* cKO and *Eogt*:*Pofut1* dKO ISC

To investigate Notch ligand binding to Notch receptors expressed in ISC, crypts were isolated, single cell suspensions prepared, and analyzed by flow cytometry. Viable, single cells were gated on CD44+CD45-CD24-ISC (Supplementary Figure S8). Cell surface expression of NOTCH1 was determined using antibody to NECD1, and binding of soluble Notch ligands carrying a C-terminal Fc tag was determined using anti-Fc antibody. Three experiments were performed, each including ISC from control, *Eogt* KO, *Pofut1* cKO and *Eogt*:*Pofut1* dKO mice. Representative flow cytometry profiles and histograms of mean fluorescence index (MFI) are shown in Figure 5. MFI data for mutant ISC were normalized to that of control ISC in each experiment. The expression of NOTCH1 on ISC cell populations from mutant mice varied but did not differ significantly among groups (Fig. 5D). Binding of DLL1-Fc was reduced compared to control in *Pofut1* cKO and *Eogt:Pofut1* dKO ISC. DLL1-Fc binding further reduced in *Eogt:Pofut1* dKO as compared to *Pofut1* cKO. Binding of DLL4-Fc was similarly reduced in mutant ISC but did not reach significance.

**Figure 5.**
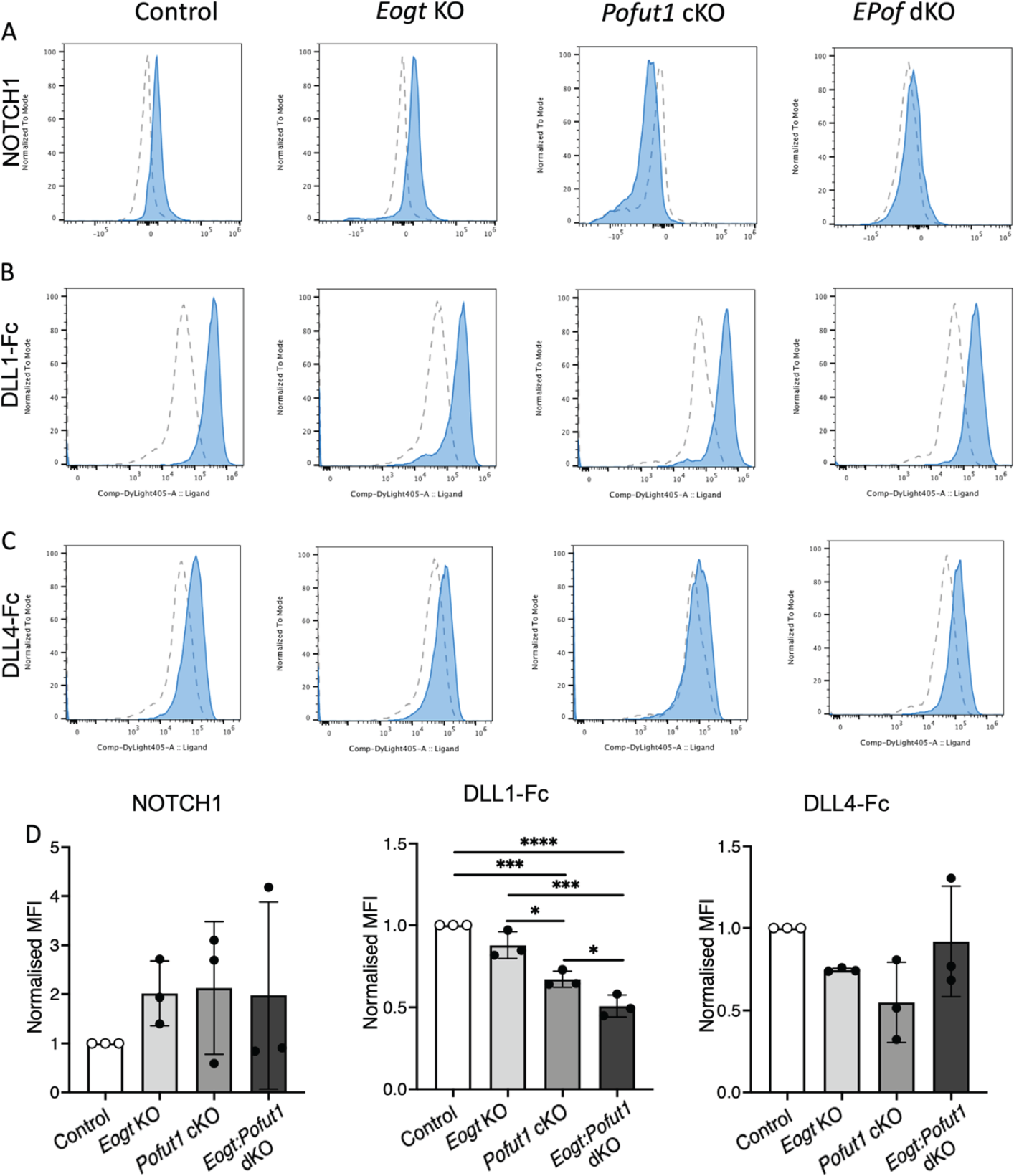
Notch ligand binding to intestinal stem cells. (*A-C*) Flow cytometry profiles of binding of anti-NOTCH1 ECD, DLL1-Fc and DLL4-Fc to *Eogt* KO, *Pofut1* KO or *Eogt:Pofut1* dKO intestinal stem cells. (*D*) MFI for anti-NOTCH1 ECD, Notch ligand DLL1-Fc and Notch ligand DLL4-Fc (n = 3 mice) binding to Control, *Eogt* KO, *Pofut1* KO and *Eogt:Pofut1* dKO normalized to MFI of Control cells. Average MFI for NOTCH1 ECD binding was Control, 3639; *Eogt* KO, 6245; *Pofut1* KO, 4623; *Eogt:Pofut1* dKO, 5837. Average MFI for DLL1-Fc binding was Control, 213,525; *Eogt* KO, 184,167; *Pofut1* KO, 143871; *Eogt:Pofut1* dKO, 105,452. Average MFI for DLL4-Fc binding was Control, 39,905, *Eogt* KO, 29,691; *Pofut1* KO, 20562; *Eogt:Pofut1* dKO, 40779. Blue profiles show NOTCH1 ECD or Notch ligand-Fc binding, gray profiles show secondary antibody binding. *P* values from one-way ANOVA followed by Tukey’s multiple comparisons test -\**P* < 0.05, \*\*\**P* < 0.001, \*\*\*\**P* < 0.0001.

### Notch Target and Notch Pathway Gene Expression in *Eogt:Pofut1* dKO Intestine

To investigate the functional impact of EOGT and POFUT1 synergism, expression of Notch signaling target genes, lineage and ISC marker genes and Notch pathway member genes were investigated for crypt cells. Fractionation produced a similar enrichment of villi versus crypt cells regardless of genotype (Supplementary Figure S2A and S2B). *Hes1* is directly regulated by Notch signaling and is decreased by reduced Notch signaling in the intestine, leading to increased expression of targets such as *Math1* ^13, 26^. Consistent with this, *Hes1* expression was markedly decreased in *Pofut1* cKO crypts. Importantly, it was further decreased in *Eogt*:*Pofut1* dKO crypts (Fig. 6A). In parallel, *Math1* expression increased in both *Pofut1* cKO and *Eogt*:*Pofut1* dKO crypts (Fig. 6A). Further, transcripts of secretory cell marker genes *ChgA* and *Lyz1* in both mutants were increased (Fig. 6B) as expected based on histological analyses that revealed increased secretory cells in *Pofut1* cKO intestine, and an even greater increase in *Eogt*:*Pofut1* dKO intestine (Fig. 3C and 3E). *Muc2* transcripts were also increased in both *Pofut1* cKO and double mutant crypts (Fig. 6B). Expression of two canonical stem cell marker genes, *Lgr5* and *Olfm4* were both decreased in single and double mutant crypts (Fig. 6C). Transcripts of *Notch1* were decreased in both *Pofut1* cKO and *Eogt:Pofut1* dKO crypts, while *Notch2* expression was decreased in only *Eogt*:*Pofut1* dKO crypts (Fig. 6D). *Jag1* expression was also decreased in both mutants (Fig. 6D). In contrast, *Dll1* expression was increased in *Eogt*:*Pofut1* dKO crypts while *Dll4* expression was increased in both mutants. *Jag2* transcripts were also increased in both of the mutants compared to controls, and increased to a greater extent in *Eogt*:*Pofut1* dKO compared to *Pofut1* cKO crypts (Fig. 6D). Overall, therefore, the expression of marker genes of Notch signaling support the hypothesis that EOGT contributes to canonical, Notch ligand-mediated signaling in maintaining homeostasis of the intestinal mucosa.

**Figure 6.**
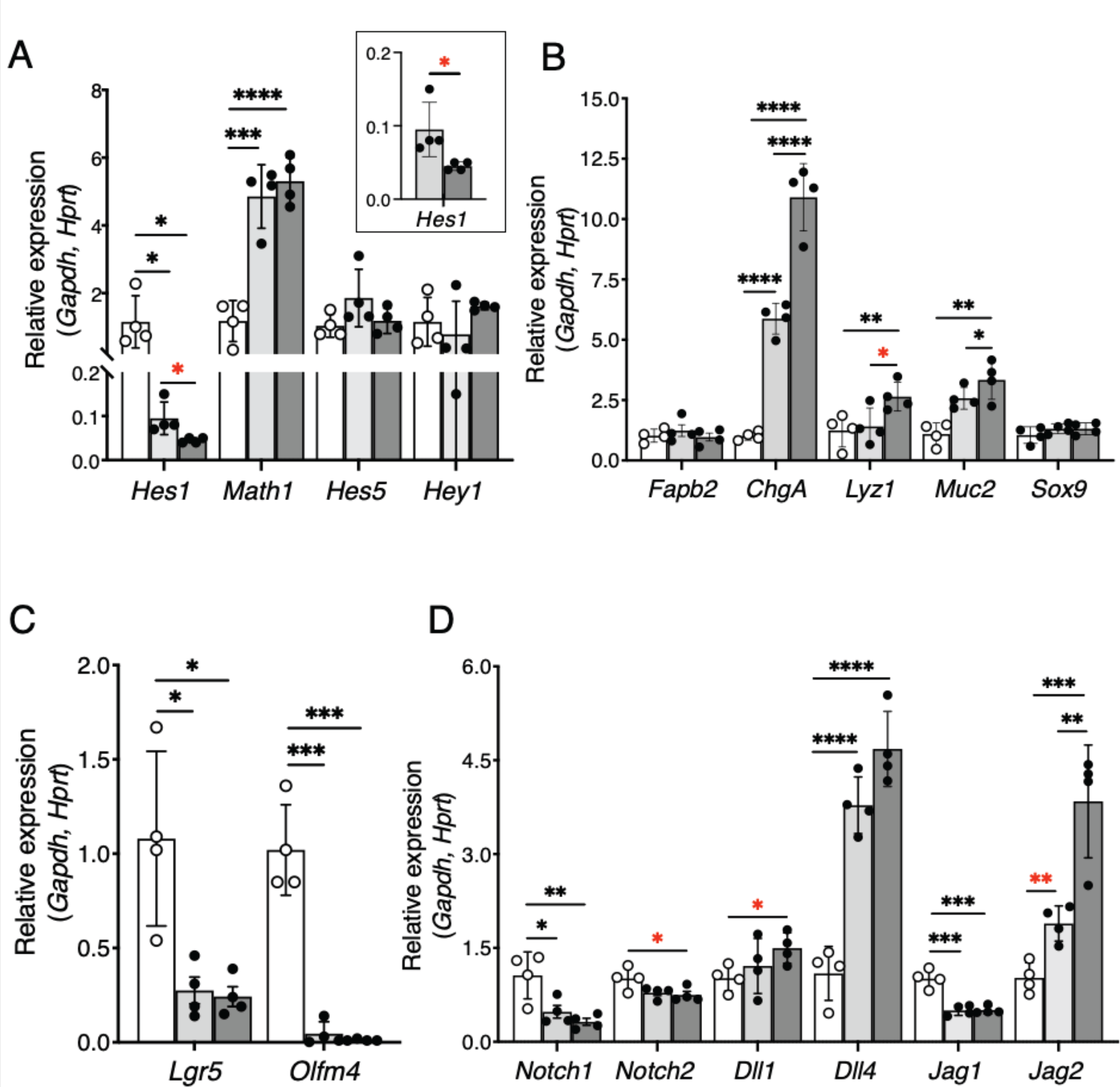
Notch pathway genes expressed in crypts of *Pofut1* cKO and *Eogt:Pofut1* dKO mice. (*A*) qRT-PCR performed on cDNA from intestinal crypts. Transcript levels of key Notch pathway target genes in Control (white), *Pofut1* cKO (light gray) and *Eogt:Pofut1* dKO (dark gray) crypt cells. Transcript levels of marker genes expressed in (*B*) different intestinal cell lineages, or (*C*) ISC. (*D*) Transcript levels of Notch receptors and Notch ligands (n = 4 mice per group). P values from one-way ANOVA followed by Tukey’s multiple comparisons test - *P < 0.05, **P < 0.01, ***P < 0.001, ****P < 0.0001, or student’s t-test with Welch’s correction *P < 0.05, **P < 0.01

## Discussion

Here we show that Notch signaling in mouse intestine is fine-tuned by the addition of two types of O-glycan to Notch receptors that modulate the binding of canonical Notch ligands - O-fucose and O-GlcNAc glycans. While the contribution to Notch signaling strength of POFUT1 and O-fucose glycans is much stronger than that of EOGT and O-GlcNAc glycans, the necessity for the latter became clear in *Eogt:Pofut1* dKO intestine. Whereas *Pofut1*:Villin1-Cre cKO mice exhibit severely defective Notch signaling in mouse intestine ^13^ (and herein), *Pofut1* cKO mice live for ≥ 6 months, whereas we establish here that *Eogt:Pofut1*:Villin1-Cre dKO mice die at ∼1 month with a more severely defective Notch signaling intestinal phenotype. Nevertheless, these compound mutants live longer than mice lacking both *Notch1* and *Notch2* or *Dll1* and *Dll4* which die within days of conditional knockout ^15, 16^. We propose that limited Notch signaling persists in *Eogt:Pofut1* dKO intestine via Notch receptors that are modified by only O-glucose glycans (Fig. 1A). While O-glucose glycans do not appear to interact directly with Notch ligands ^10, 11, 22^, they are important in regulating Notch receptor trafficking to the cell surface ^27, 28^. Since anti-NOTCH1 ECD antibodies and soluble Notch ligands DLL1-Fc and DLL4-Fc bound to ISC from *Eogt:Pofut1* dKO intestine (Fig. 5), Notch receptors had clearly trafficked to the cell surface. We propose that O-GlcNAc glycans on Notch receptors support that trafficking, and that elimination of Notch signaling by reducing glycosylation would require the triple knockout of *Pofut1, Eogt* and *Poglut1*.

Revealing that *Eogt* contributes to Notch signaling in mouse intestinal development is important for several reasons. First, to better understand mechanistic bases of regulation of the ubiquitously important Notch signaling pathway. Second, to understand potential pathologies in individuals with the congenital disease of glycosylation EOGT-CDG (Adams-Oliver Syndrome Type 4; AOS4). Those afflicted lack a functional *EOGT* gene and exhibit a range of limb extremity and scalp developmental defects as well as ocular, vascular and intellectual abnormalities consistent with defective Notch signaling ^29, 30^. In the mouse embryo, the *Eogt* gene is first expressed in limb buds and the apical ectodermal ridge before expression in digit condensates at ∼E12.5, consistent with defective development of fingers and toes in EOGT-CDG patients ^29^. The involvement of *Eogt* in mouse intestinal development reported here is a potential indication of the possibility of perturbed intestinal development or function in EOGT-CDG patients. Mutations in *EOGT* have been associated with 2% of 10,967 colorectal adenocarcinoma patients queried ^31^.

Determining how EOGT and the O-GlcNAc glycans fine tune Notch functions in the intestinal environment is challenging. Ideally, NOTCH1 from ISC from the various mouse cohorts would be purified and the nature of the attached O-glycans defined, but this is currently not feasible due to the low concentration of NOTCH1 and NOTCH2 in ISC. The first O-glycan analysis of mouse NOTCH1 from a physiologic source required stimulation of splenic T cells in culture to increase *Notch1* expression and obtain sufficient NOTCH1 for mass spectrometric analysis ^32^. Roles for the O-glycans on canonical Notch ligands must also be evaluated. Deletion of *Rfng* is reported to reduce modification of Delta ligands and their ability to stimulate Notch signaling in ISC ^14^. However, determining which under-glycosylated partner in a defective Notch receptor-ligand interaction is responsible can only be done when the other partner is fully glycosylated. Thus, proving that Notch ligands require specific O-glycans to induce Notch signaling necessitates conditionally deleting the target glycosylation gene in ligand-expressing cells. Finally, to conclude that Notch signaling is affected by the deletion of a glycosyltransferase requires evidence that the loss generates: 1) a phenotype that mimics loss of Notch pathway member(s); 2) causes alterations in Notch ligand binding; and 3) causes altered expression of Notch signaling target genes. Here we provide this combined evidence, demonstrating that EOGT and the O-GlcNAc glycans it initiates support Notch signaling during intestinal development in the absence of POFUT1 and O-fucose glycans.

## Methods

### Single cell RNAseq bioinformatics

Bioinformatic trajectory analysis of scRNAseq data obtained from Epcam+CD45-C57Bl6/J adult intestinal crypt cells and deposited as GSE188339 was performed as described ^33^

### Primers and Antibodies

Primer sequences are given in Supplementary Table 1 and antibodies are given in Supplementary Table 2.

### Generation of Chinese Hamster Ovary (CHO) Glycosylation Mutants

CHO Lec1 ^23^ cells with inactivated *Pofut1* were previously described as *Pofut1* KO-1 (P4-11) and KO-2 (P4-15) ^34^. The *Pofut1* deletion was confirmed by western blotting with a rabbit anti-bovine POFUT1 antibody ^35^. To inactivate *Eogt*, two guide RNAs (*GTATGACTACTCCAGCCTCC* in exon 1; Eogt2 *GTTTGCAGCTATGTCGACGT* in exon 2), an ATTO 550-labeled tracrRNA and Cas9 (all from Integrated DNA Technologies, Inc., Coralville, IA) were lipofected into Lec1 cells. After 24 hours, ATTO 550+ cells were sorted by flow cytometry and the 3% most positive cells were seeded at 1 cell per well in alpha minimal essential medium (αMEM, Thermo Fisher Scientific, Waltham, MA) containing 10% fetal bovine serum (FBS) and 1% Penicillin-Streptomycin. Lec1 *Pofut1* KO-1 cells were transformed by lipofection with the same *Eogt* gRNAs to generate *Eogt:Pofut1* dKO mutants. After 8-10 days, colonies were expanded and characterized. Genomic DNA PCR detected *Eogt* alleles (Supplementary Figure S1A) and RT-PCR detected cDNA products (Supplementary Figure S1B) as described ^19^. Mutants were designated E116 (*Eogt* KO) and PE316 (*Eogt:Pofut1* dKO).

### Notch Ligand Binding to CHO Mutants

Cells (∼0.5 × 10^6^) were washed with FACS binding buffer (FBB; Hank’s balanced salt solution (MilliporeSigma Corning, Burlington, MA), 2 % bovine serum albumin (BSA) fraction V (GeminiBio, West Sacramento, CA), 1 mM CaCl_2_ and 0.1 % sodium azide) and incubated in 50 μl of rat anti-mouse CD16/CD32 Fc blocker (Supplementary Table 2 for all antibodies) in FBB for 15 minutes on ice. Sheep anti-mouse NOTCH1 (100 μl diluted 1:50 in FBB) or 100 μl FBB containing 0.75 μg DLL1-Fc, DLL4-Fc or JAG1-Fc soluble Notch ligand described previously ^35^ was added to the Fc block and incubated on ice for 1 hour, washed twice with cold FBB and incubated with secondary antibody in FBB (1:100 rhodamine Red-X anti-sheep IgG) or anti-Fc for ligands (1:100, Alexa Flour 488 anti-human IgG) for 30 minutes on ice. Secondary antibody alone determined background binding. Following addition of 5 μl 7-AAD in 95 μl FBB for 10 minutes at room temperature, FBB (250 μl) was added, cells filtered into 5 ml polystyrene tubes and analyzed by flow cytometry using a FACS Caliber flow cytometer and FlowJo software (BD Biosciences, Franklin Lakes, NJ).

### Mouse Models

Mice with a targeted inactivating mutation in the *Eogt* gene were developed at Nagoya University and previously described ^19^. *Pofut1*[F/F] mice were also generated previously ^36^. Mice expressing Villin-Cre (B6.Cg-Tg(Vil1-cre)1000Gum/J, JAX strain 021504) were from Jackson Laboratories, Portland, MA. Cross breeding generated *Pofut1* cKO (*Eogt*[+/-]:*Pofut1*[F/F]:Villin-Cre), *Pofut1*:*Eogt* dKO (*Eogt*[-/-]*Pofut1*[F/F]:Villin-Cre) and mutants lacking Villin-Cre. Genomic DNA PCR was used to genotype. Body weight was recorded following euthanasia, small intestine was dissected, length measured, and tissue processed. Experimental protocols were approved by the Albert Einstein Institutional Animal Care and Use Committee under protocol numbers 20170709 and 00001311.

### Isolation of Crypt and Villus Fractions

Small intestine was washed with cold phosphate-buffered saline containing 1 mM CaCl_2_, 1 mM MgCl_2_ and 1 mM dithiothreitol (PBS (Ca/Mg/DTT), sectioned into four, flushed extensively with PBS (Ca/Mg/DTT), one end tied, everted with a gavage needle, filled to ∼75% with PBS (Ca/Mg/DTT) and tied off before washing with buffer A (96 mM NaCl, 1.5 mM KCl, 27 mM Na-Citrate, 8 mM KH_2_PO_4_, 5.6 mM Na_2_HPO_4_ and 1 mM DTT) at 37 °C with constant shaking for 10 minutes. Tissue was shaken with buffer B (1.5 mM EDTA, 0.5 mM DTT and 0.1 % BSA) four times at 37 °C for 10, 10, 30 and 20 minutes, respectively. Buffer B collected at each time was centrifuged to give fractions I-IV. Relative enrichment of villi and crypts was determined by qRT-PCR of marker genes (Supplementary Figure S2A and S2B). Fractions were snap frozen in liquid N_2_ and stored at -80 °C. Conditional deletion of *Pofut1* by Villin-Cre was validated by PCR genotyping (Supplementary Figure S2C).

### Histopathology and Immunohistochemistry

Jejunum was fixed in 10% neutral buffered formalin for 48 hours and processed through a graded series of alcohol to prepare paraffin blocks. Sections (5 μm) were stained with hematoxylin and eosin or Alcian Blue. Slides were scanned using a 3D Histech P250 High-Capacity Slide Scanner. CaseViewer 2.4 was used to measure villi length, crypt depth, crypt width and crypt length. QuantCenter software was used to quantitate goblet and Paneth cell staining. For immunohistochemistry, 5 μm sections were dipped into xylene and graded concentrations of ethanol as described (http://www.abcam.com/protocols/). Slides were boiled in sodium citrate for 20 minutes, incubated with 0.1% Triton X-100 (only for cleaved anti-NICD1) followed by 1.5% H_2_O_2_ in Tris-buffered saline (TBS) for 20 minutes. Antibody incubations were performed at room temperature in a humidified chamber as follows: blocking in 10% FBS with 1% BSA in TBS for 1 hour, followed by primary antibody overnight in 1% BSA in TBS, washing with TBS containing 0.025% Triton X-100, and incubation in HRP-conjugated secondary antibody diluted in TBS with 1% BSA for 1 hour. Diaminobenzidine peroxidase substrate kit (Vector Laboratories, Burlingame, CA) treatment was followed by counter staining with Hematoxylin and Bluing reagent. Slides were treated with a graded ethanol series and xylene and mounted using Permount.

### Quantitative RT-PCR (qRT-PCR)

Total RNA was extracted from ∼10^7^ frozen cells using TRIzol (Thermo Fisher Scientific, Waltham, MA). RNA was dissolved in RNAse-free water and 1 µg estimated using Nanodrop was used to make 20 µl cDNA (Verso cDNA synthesis kit, Thermo Fisher Scientific). cDNA was amplified using the PowerUp SYBR Green master mix (Thermo Fisher Scientific) and 750 nM each primer. Vii7 Real-Time PCR system (Applied Biosystems, Foster City, CA) was used to run qRT-PCR for 40 cycles. Each sample was run in triplicates using 384 wells plate. Gene expression was calculated relative to *Gapdh* and *Hprt* by log2^ddCT^ method.

### Western Blotting

Frozen cells (∼10^7^) were homogenized in 100 μl lysis buffer containing 1% IGEPAL, 1%TX-100, 0.5% Deoxycholate (all from Sigma-Aldrich, St. Louis, MO), and Roche complete*™* Protease Inhibitor (Sigma-Aldrich) and incubated on ice for 30 minutes. After centrifugation at 5000 *g* for 5 minutes at room temperature, the supernatant in a new tube received 20% glycerol and protein concentration was estimated by Bradford’s Dye Reagent Concentrate (BioRad Laboratories, Hercules, CA). For gel electrophoresis, 50-100 µg protein was analyzed by SDS-PAGE, transferred to PVDF membrane and blocked in 5% Non-fat dry milk in Tris-buffered saline 0.05% Tween 20 followed by overnight incubation with primary antibody in blocking buffer at 4°C. Membrane rinsed with Tris-saline was incubated with HRP-secondary antibody in the same buffer for one hour at room temperature. Enhanced chemiluminescent substrate was used and signals visualized on X-ray film (Thermo Fisher Scientific).

### Notch Ligand Binding to ISC

Washed small intestine pieces were scraped to remove villi, transferred to 20 ml ice-cold 2 mM EDTA and 2 mM glutamine in PBS (Ca/Mg free) and incubated on ice for 20 minutes. Tissue was then transferred to fresh 20 ml PBS (Ca/Mg free) with 2 mM glutamine and shaken by hand for 30 seconds. This process was repeated another four times. The last four washes were filtered through a 70 μm strainer into 1% BSA/PBS (Ca/Mg free)-coated 50 ml falcon tubes. Filtrates were spun at 1500 rpm for 10 minutes at 4°C to pellet crypts. Single cells were prepared in 3 ml enzyme-free dissociation buffer (Gibco, Thermo Fisher Scientific, Waltham, MA) incubated in a 37 °C water bath for 10 minutes, with pipetting every 30 seconds, then 7 ml of alpha MEM was added, cells were filtered through a 40 μm strainer, pelleted and resuspended in Zombie NIR dye in PBS (1:7000; Biolegend). After 30 minutes rocking at 4°C, cells were washed with PBS, fixed in 4% paraformaldehyde (PFA; Emsdiasum, Hatfield, PA) in PBS (Ca/Mg free) 15 minutes at 4°C with rocking, washed 3 times with cold FBB and stored in FBB at 4°C.

Flow cytometry performed within 1-6 days investigated NOTCH1 expression using anti-NECD1 antibody AF5267 and Notch ligand binding for DLL1-Fc (R&D Systems, Inc., Minneapolis, MN) and DLL4-Fc (Aro Biosystems, Newark, DE). Briefly, ∼10^6^ fixed cells were resuspended in 40 μl CD16/CD32 Fc blocker in FBB (1:40) and incubated 15 minutes on ice. Antibodies in FBB to CD45 (1:400), CD44 (1:800), CD24 (1:800), CD166 (1:800) and GRP78 (1:800) with anti-NOTCH1 or 2 μg of Notch ligand-Fc were added to cells in FcR block and incubated 30 minutes on ice. Cells with no anti-NOTCH1 or ligand-Fc determined background. Cells were washed and incubated with secondary antibody (1:600 rhodamine Red-X conjugated donkey anti-sheep IgG or 1:100 Fc-specific anti-IgG-Dylight 405) for 30 minutes on ice, washed, resuspended in 300 μl FBB, filtered into flow tubes and 300,000 events were recorded in a CytekTM Aurora flow cytometer, analysis by FlowJo software (BD Biosciences).

## Statistical Analysis

GraphPad Prism 9.0.1 was used to perform one-way ANOVA followed by Tukey’s multiple comparisons tests or unpaired, two-tailed Student’s t tests as noted. Data are represented as mean ± SEM.

## Acknowledgements

We thank the Histology and Comparative Pathology Core, Albert Einstein College of Medicine, for sectioning and staining fixed intestine samples. We thank Andrea Briceno and Hillary Guzik, Analytical Imaging Facility, Albert Einstein College of Medicine, for scanning histology slides using a 3D Histech P250 High-Capacity Slide Scanner purchased with NIH SIG #1S10OD019961-01. We are grateful to Jinghang Zhang and Aodengtuya of the Flow Cytometry Core, Albert Einstein College, for support in designing and executing flow cytometry experiments. We also thank Subha Sundaram for overall technical assistance.

## Author Contributions

Conceptualization, PS; Methodology, NM, SV, PS, JC, LHA; Investigation, NM, SV; Analysis, NM, SV, JC, LHA, PS; Resources, PS, LHA; Writing first draft, NM, PS; Review and editing, PS, NM, LHA, JC; Visualization, NM, JC; Funding acquisition, PS, LHA.

## Data availability

The data presented in this paper will be made available upon request.

## Additional information

The authors have no conflicts of interest to disclose in relation to this work. This work was funded by National Institutes of Health grants R01 GM106417 (PS), R01 CA229216 (LHA), and partially supported by the Albert Einstein Cancer center grant PO1 CA13330.

## Abbreviations

BSA, bovine serum albumin; CHO, Chinese hamster ovary; cKO, conditional knockout; DAB, 3,3′-diaminobenzidine; Div, dividing; dKO, double knockout; DTT, dithiothreitol; EC, enterocyte; EDTA, ethylenediamine tetra-acetic acid; EE, enteroendocrine; EGF, epidermal growth factor-like; FBS, fetal bovine serum; gRNA or crRNA, guide RNA; ISC, intestinal stem cell; NECD1, NOTCH1 extracellular domain; NICD1, NOTCH1 intracellular domain; R, replicating; RNP, ribonucleoprotein; sc, single cell; TA, transit amplifying; UK, unknown.

