## Supplemental material for "EOGT Enables Residual Notch Signaling in Mouse Intestinal Cells Lacking POFUT1"

**Short Title:** Notch O-glycans act synergistically in mouse intestine

Mohd Nauman [1], Shweta Varshney [1,2] Jiahn Choi [1,3], Leonard H. Augenlicht [1,3] and Pamela Stanley [1],*

[1] Dept. Cell Biology, Albert Einstein College of Medicine, New York, NY 10461

[2] Current address: Dudnyk, 5 Walnut Grove Drive, Suite 300, Horsham, PA 19044

[3] Depts. Medicine and Oncology, Albert Einstein College of Medicine, New York, NY 10461

**Supplementary Files – Table of Contents**

1. **Supplementary Tables**

    Supplementary Table 1

    Supplementary Table 2

2. **Supplementary Figures**

    Supplementary Figure 1

    Supplementary Figure 2

    Supplementary Figure 3

    Supplementary Figure 4

    Supplementary Figure 5

    Supplementary Figure 6

    Supplementary Figure 7

    Supplementary Figure 8

**Supplementary Table 1.** Primers used in this study.

| Gene | Name | Sequence 5'-3' |
|---|---|---|
| **CHO DNA, cDNA** | | |
| ***Eogt*** | F (2F) | *CCACTGACCTACATGCCCACCCTTAAA* |
| | R (10R) | *CAACAGAAACCTATGAAGTTAGGGTTGTCTGG* |
| ***Eogt*** | F (SV20F) | *ATGTTAATGCTGCTTGCCTTTGGAG* |
| | R (SV20R) | *CTGAAAGAATTGTTCACGTGCTGGG* |
| **Mouse DNA** | | |
| ***Pofut1 F/del*** | F (PS644) | *GGGTCACCTTCATGTACAAGTGAGTG* |
| | R (PS645) | *ACCCACAGGCTGTGCAGTCTTTG* |
| ***Pofut1 F/WT*** | F (FB21) | *CCAGGCTGATCACTTCTTGG* |
| | R (FB22) | *CCCTGTCTCGAAAAAGCAAA* |
| ***Eogt*** | F (3rd loxF) | *CCACCCGACCCCTGCCAGAACATAATGCTCTCTTGCATC* |
| | R (3rd loxR) | *GCTGTCGCCAGAGGAGAGAGTGGGTGCTTACTTAC* |
| | R (25307 Rv) | *CCAAGGCGGTCTTGGCCCAT* |
| ***Villin-Cre*** | F (16775) | *GCCTTCTCCTCTAGGCTCGT* |
| | R (16776) | *TATAGGGCAGAGCTGGAGGA* |
| | R (oIMR9074) | *AGGCAAATTTTGGTGTACGG* |
| **qRT-PCR** | | |
| ***Hes1*** | F | *AGCTGGAGAGGCTGCCAAGGTTT* |
| | R | *ACATGGAGTCCGAAGTGAGCGAG* |
| ***Hes5*** | F | *GGACCAGAGGATGAGCTCGTT* |
| | R | *AGGAGGGAGCCTTCGGAAGA* |
| ***Hey1*** | F | *TGAGCTGAGA AGGCTGGTAC* |
| | R | *ACCCCAAACTCCGATAGTC* |
| ***Math1*** | F | *ATGCACGGGCTGAACCA* |
| | R | *TCGTTGTTGAAGGACGGGATA* |
| ***Notch1*** | F | *GAGATGCCCTCGGACCAATC* |
| | R | *GGCGAAAGCGAGAACACA* |
| ***Notch2*** | F | *TGTACCAGATCCCAGAGATGC* |
| | R | *GTCAGATGCAGAGTGTGGTGA* |
| ***Dll1*** | F | *GTCTGCCAGGGTGTGATGAC* |
| | R | *CGGATGCACTCATCGCAGTA* |
| ***Dll4*** | F | *AGTGCCAGAACAGAGGTCCAA* |
| | R | *CAGGGACTTCGGGCACAAT* |

| | | |
|---|---|---|
| ***Jag1*** | F<br>R | *TGACATGGATAAACACCAGCA*<br>*GCAGCCCACTGTCTGCTATAC* |
| ***Jag2*** | F<br>R | *ATTGTAGCAAGGTATGGTGCG*<br>*GCACAGTTGTTGTCCAAATGA* |
| ***Lgr5*** | F<br>R | *TCTCCTACATCGCCTCTGCT*<br>*TTCCTCCGGAACCTGTCTCA* |
| ***Olfm4*** | F<br>R | *ATTCGCTATGGCCAAGGAGG*<br>*GAGGGGCCGATTCACATCAA* |
| ***Chga*** | F<br>R | *GAAGTGCGCCTGGAAGTCA*<br>*GATCCTCTCGTCTCCTTGGA* |
| ***Fabp2*** | F<br>R | *CTAGAGACACACACAGCTGAGATCATGG*<br>*GCAATCAGCTCCTTTCCATTGTCTACAC* |
| ***Hprt*** | F<br>R | *TCAGTCAACGGGGGACATAAA*<br>*GGGGCTGTACTGCTTAACCAG* |
| ***Gapdh*** | F<br>R | *GTGTCCGTCGTGGATCTGA*<br>*CCTGCTTCACCACCTTCTTG* |

F = Forward, R = Reverse

**Supplementary Table 2.** Antibodies used in this study.

| Antibody | Fluorochrome | Isotype | Clone | Cat. no. | Company |
|---|---|---|---|---|---|
| **NOTCH1** | - | Ag-purified polyclonal sheep | aa19-526 | AF5267 | R&D Systems Inc., Minneapolis, MN |
| **Cleaved NOTCH1** | - | Rabbit IgG | | 4147 | Cell Signaling Tech., Danvers, MA |
| **Cleaved NOTCH2** | - | Rabbit IgG | Polyclonal | 93-7 | G. A. Weinmaster, UCLA |
| **CD45** | PerCP | Rabbit IgG2b, κ | 30-F11 | 103129 | Biolegend, San Diego, CA |
| **CD44** | PE-Cyanine 7 | Rabbit IgG2b, κ | IM7 | 25-0441-81 | eBioscience, San Diego, CA |
| **CD24** | BV510 | Rabbit IgG2b, κ | M1/69 | | Biolegend, San Diego, CA |
| **CD166** | Alexa Fluor 700 | Goat IgG | Polyclonal | FAB1172N | R&D Systems Inc., Minneapolis, MN |
| **GRP78** | Alexa Fluor 647 | Rabbit IgG | Polyclonal | PA1-014A-A647 | Invitrogen, Carlsbad, CA |
| **Anti-sheep IgG** | Rhodamine Red-X | Donkey anti-sheep IgG | Polyclonal | 713-295-147 or 713-295-003 | Jackson ImmunoResearch, Lab. Inc., West Grove, PA |
| **Anti-human Fcγ** | Dylight™ 405 | Goat anti-human IgG | Polyclonal | 109-476-170 | Jackson ImmunoResearch, Lab. Inc., West Grove, PA |
| **Anti-human Fcγ** | Alexa Fluor 488 | Rabbit anti-human IgG | Polyclonal | A-21220 | Molecular Probes, Eugene, OR |

Supplementary Figure 1

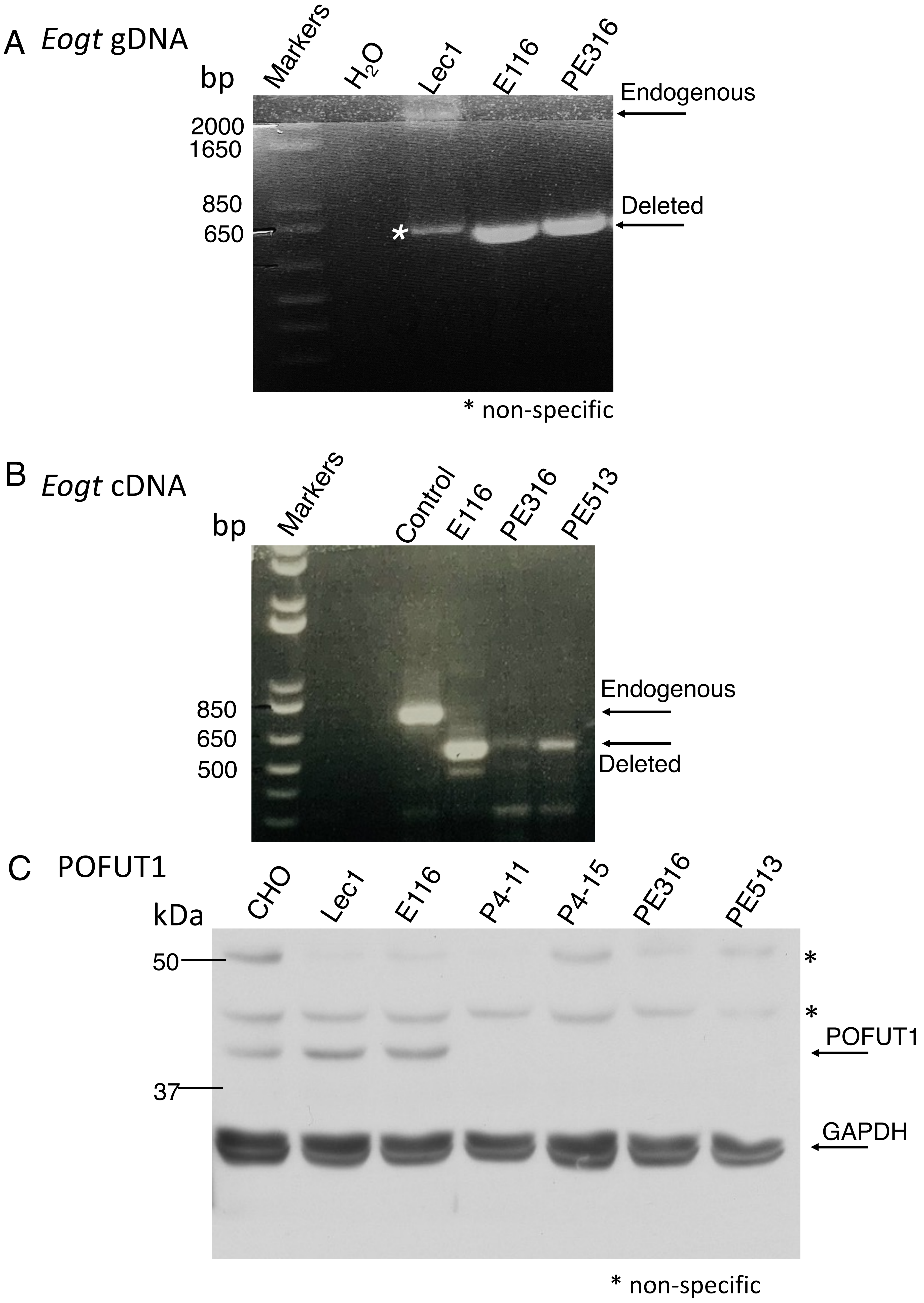

**Supplementary Figure 1.** Mutation of *Eogt* and *Pofut1* in Lec1 CHO cells. *(A)* PCR of genomic DNA (gDNA) to detect *Eogt* in E116 and PE316 cells. Endogenous *Eogt* in Lec1 cells (2198 bp) and the deleted band in E116 and PE316 cells (818 bp) are indicated. *(B)* RT-PCR of cDNA to detect endogenous *Eogt* cDNA (800 bp) in Lec1 cells and deleted band (614 bp) in E116 and PE316 cells. *(C)* Western blot analysis showing POFUT1 expression in CHO, Lec1 and E116 cells and no band for POFUT1 in P4-11, P4-15, PE316 and PE513 cells.

Supplementary Figure 2

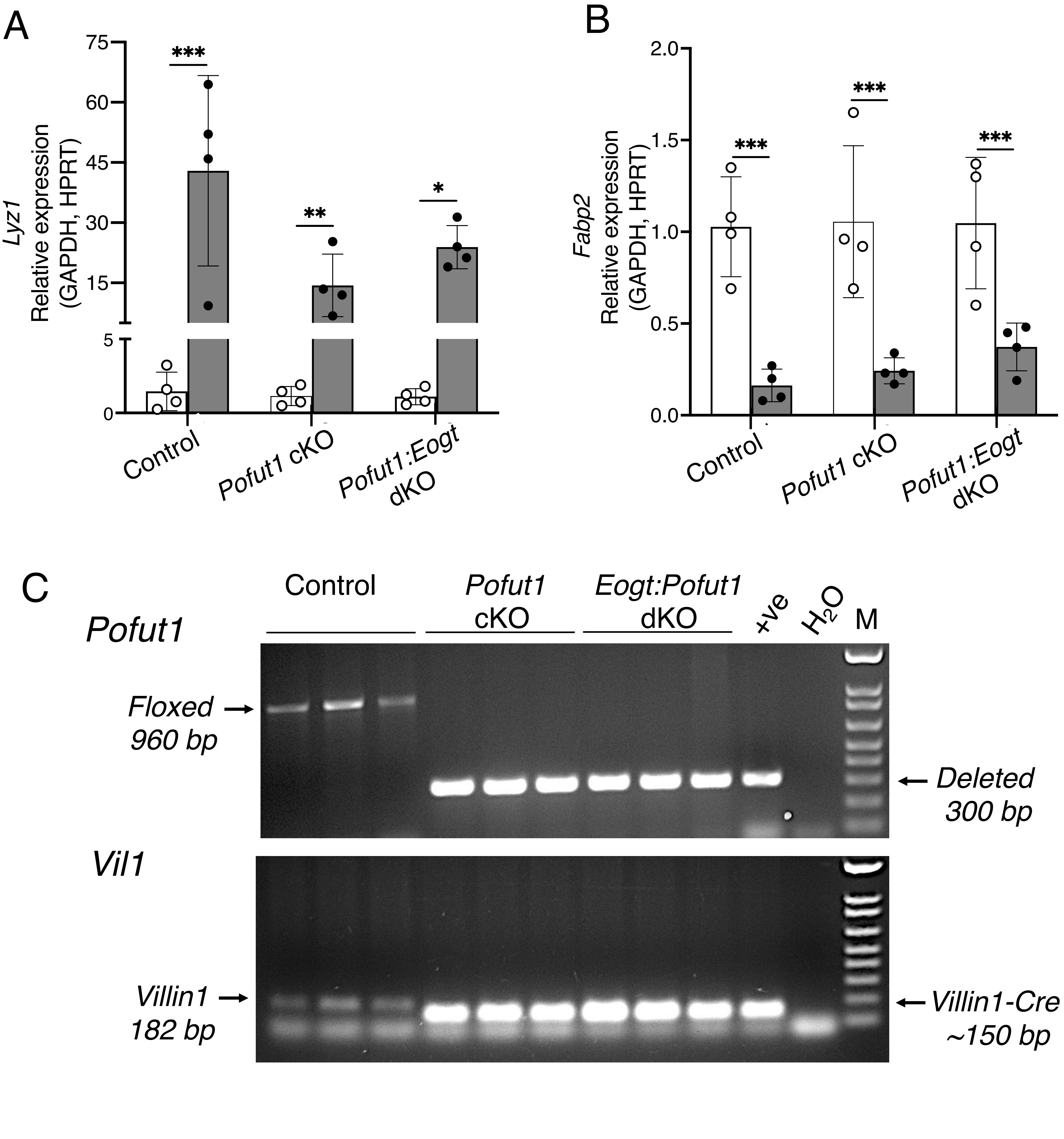

**Supplementary Figure 2.** Validation of isolated villi and crypts fractionation scheme. (*A* and *B*) Enrichment of *Lyz1* in crypts (gray) and *Fabp2* in villi (white) are shown in Control, *Pofut1* cKO and *Eogt:Pofut1* dKO fractions (n ≥ 4 mice per group). P values were determined by one-way ANOVA with Tukey's correction - *P < 0.05, **P < 0.01, ***P < 0.001 ****P < 0.0001. (*C*) Villin1-Cre deletion of *Pofut1* (300 bp) is shown in gDNA isolated from crypts of *Pofut1* cKO and *Eogt:Pofut1* dKO mice, and *Pofut1* floxed band (960 bp) is shown in gDNA from crypts of control mice (n = 3 mice per group). Controls were a known positive (+ve) and $H_2O$. M denotes 100 bp markers. The Villin1 endogenous gene (182 bp) and Villin-Cre transgene (~150 bp) were detected in Control versus *Pofut1* cKO and *Eogt:Pofut1* dKO mice (n = 3 mice per group). The endogenous gene was difficult to detect in heterozygous transgenic mice due to the robust production of the smaller PCR product from the transgene.

Supplementary Figure 3

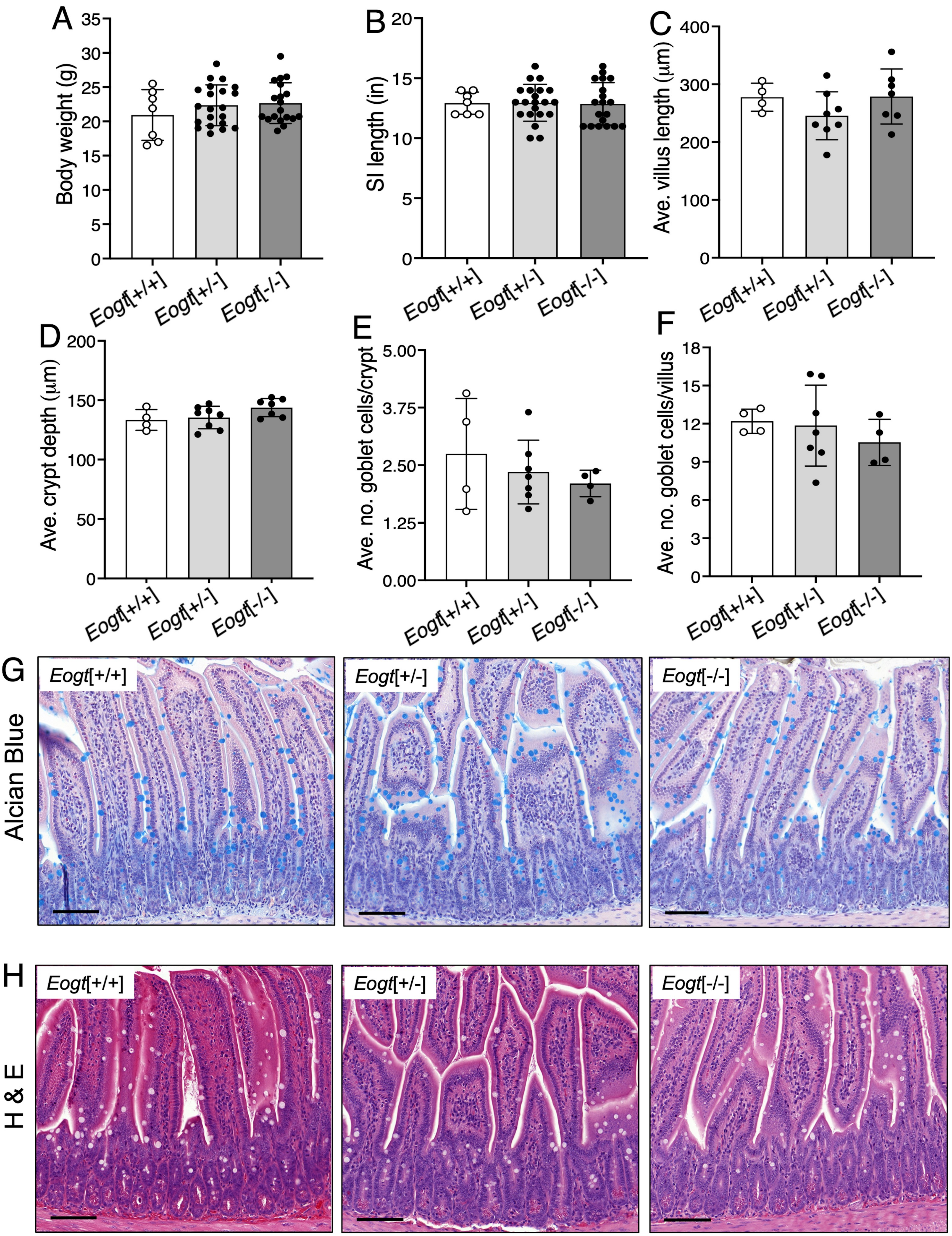

**Supplementary Figure 3.** *Eogt* deletion has no apparent effects on overall differentiation in the intestinal epithelium. (*A-F*) Comparison between *Eogt*[+/+], *Eogt*[+/-] and *Eogt*[-/-] of body weight (n ≥ 7 mice per group), length of small intestine (n ≥ 7 mice per group), villi length (20 villi were analyzed in 4 mice per group), crypt depth (20 crypts were analyzed in 4 mice per group) and, number of goblet cells in villi and crypts (30 villi and 100 crypts were analyzed in 4 mice per group). (*G* and *H*). Representative images (n = 4 mice per group) showing goblet and Paneth cells in small intestine.

Supplementary Figure 4

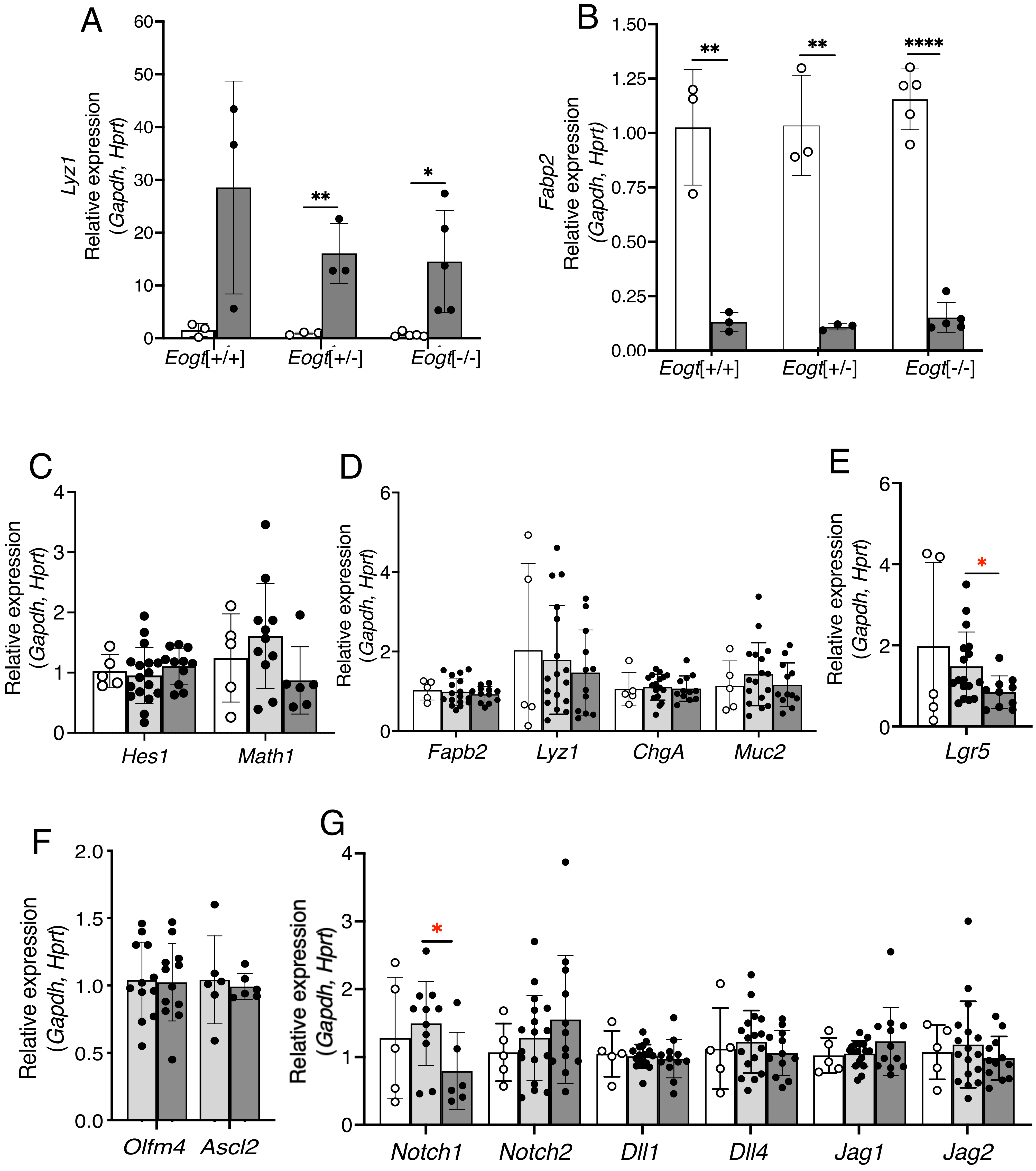

**Supplementary Figure 4.** Validation of villi and crypts fractionation and expression of Notch target and pathway genes in *Eogt* mutant mice. (*A* and *B*) Enrichment of *Lyz1* in crypts (gray) and *Fabp2* in villi (white) is shown for *Eogt*[+/+], *Eogt*[+/-] and *Eogt*[-/-] (n $\geq$ 3 mice per group). (*C*-*G*) Quantitative RT-PCR analysis of Notch signaling related genes in the small intestine in mice with partial or full deletion of *Eogt*. Transcript levels of Notch targets, markers for absorptive and secretory lineages, ISC marker(s), Notch receptors and Notch ligands in *Eogt*[+/+] (white), *Eogt*[+/-] (light gray) and *Eogt*[-/-] (light gray) (n $\geq$ 5 mice per group). P values were determined by one-way ANOVA with Tukey's correction *P < 0.05, **P < 0.01, ****P < 0.0001 or student's t test with Welch's correction - *P < 0.05.

Supplementary Figure 5

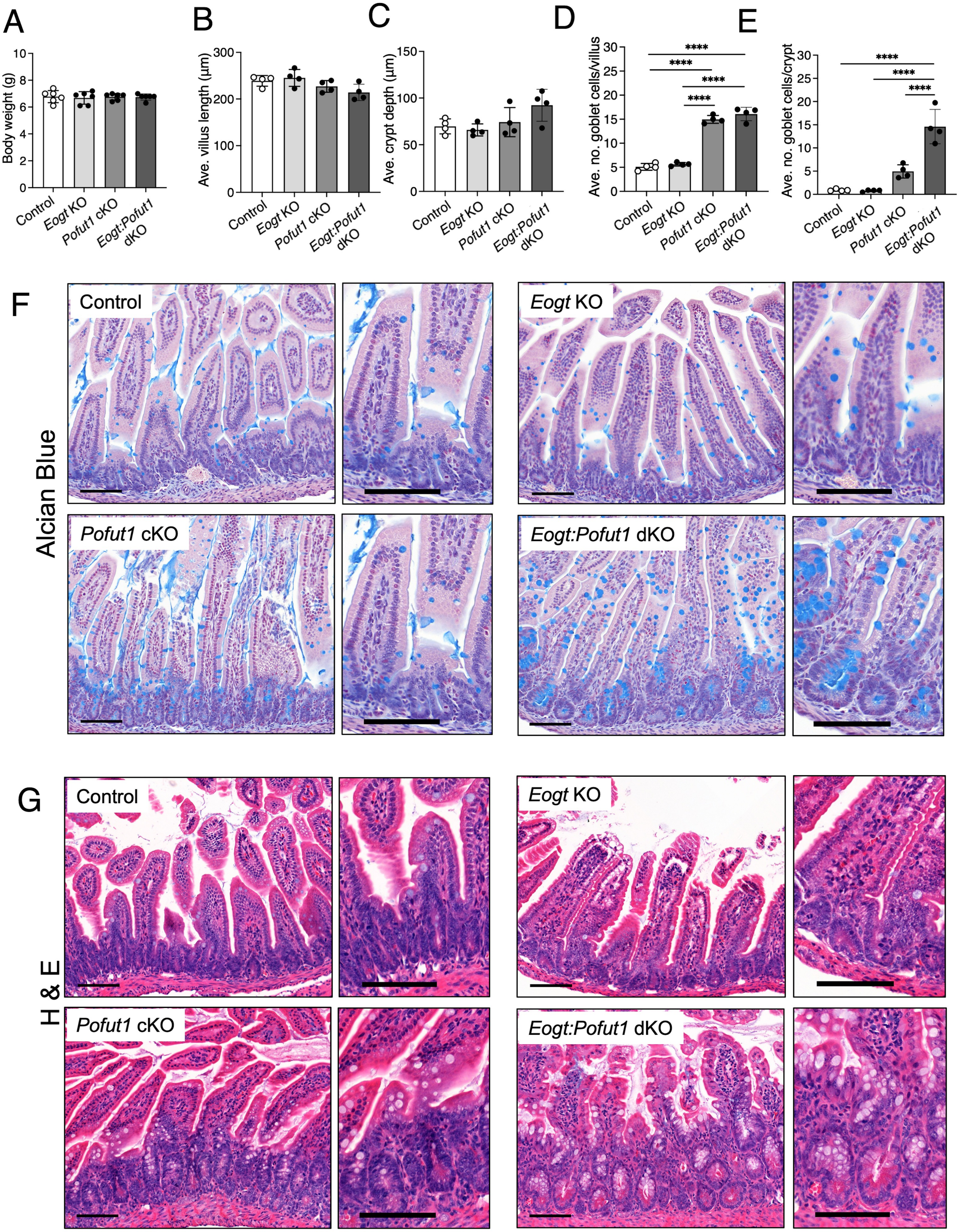

**Supplementary Figure 5.** Effects of combined deletion of *Pofut1* and *Eogt* on small intestine at P15. (*A*-*C*) Body weight (n = 6 mice per group), villus length and crypt depth of Control, *Eogt* KO, *Pofut1* cKO and *Eogt:Pofut1* dKO (20 villi or 20 crypts were analyzed in 4 mice per group). (*D* and *E*) Number of goblet cells in villi and crypts of experimental groups (30 villi and 100 crypts were analyzed in 4 mice per group). (F and G) Representative images showing goblet and Paneth cells in the small intestine of experimental mice (n = 4 mice per group). P values were determined by one-way ANOVA with Tukey's correction - ****P < 0.0001.

Supplementary Figure 6

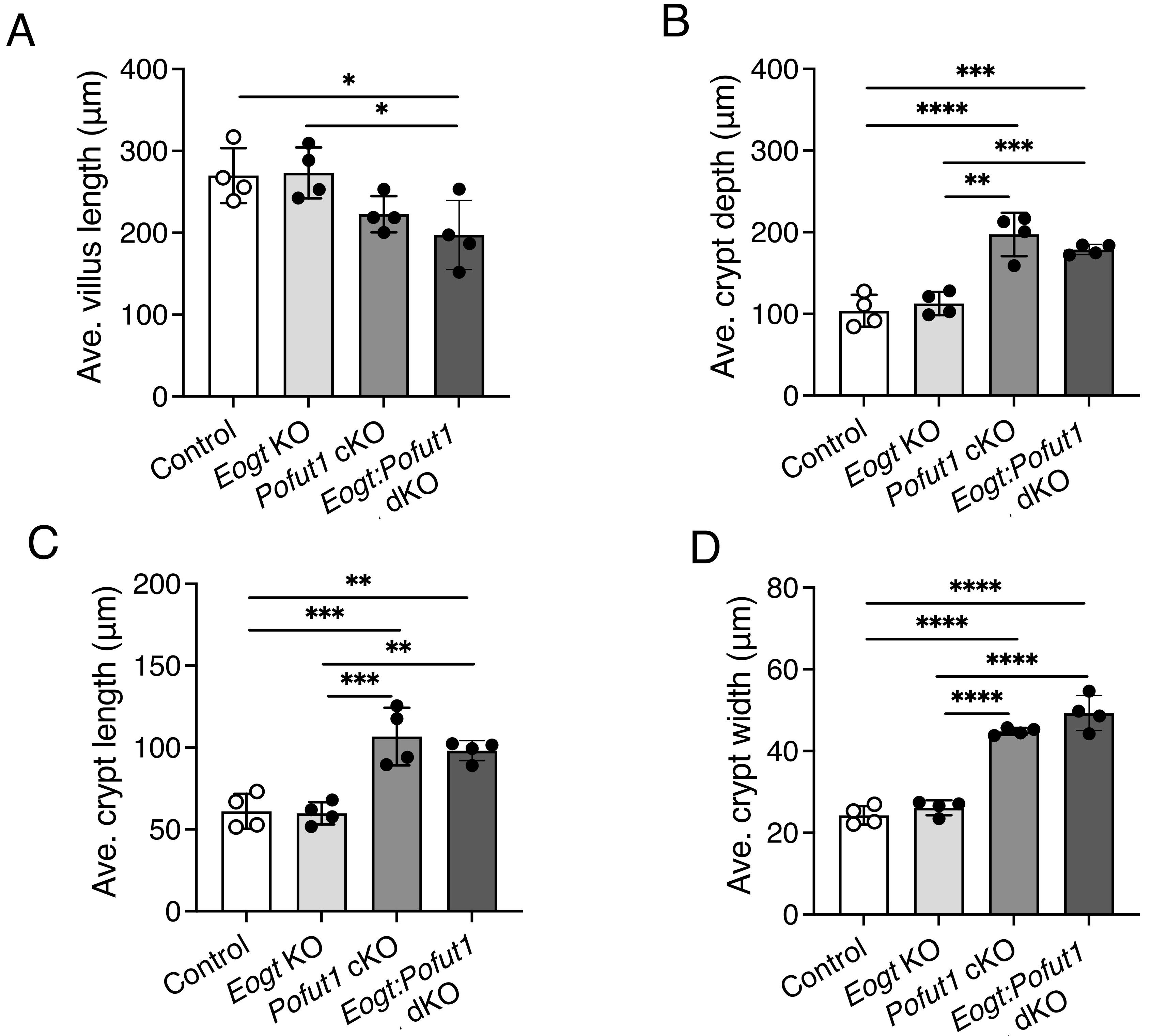

**Supplementary Figure 6.** Effects of *Pofut1* deletion with and without *Eogt* at P28. (*A-D*) Comparisons of villi length, crypt depth, crypt length and crypt width in Control, *Eogt* KO, *Pofut1* cKO and *Eogt:Pofut1* dKO (n = 4 mice per group). P values were determined by one-way ANOVA with Tukey's correction - *P < 0.05, **P < 0.01, ***P < 0.001, ****P < 0.0001.

Supplementary Figure 7

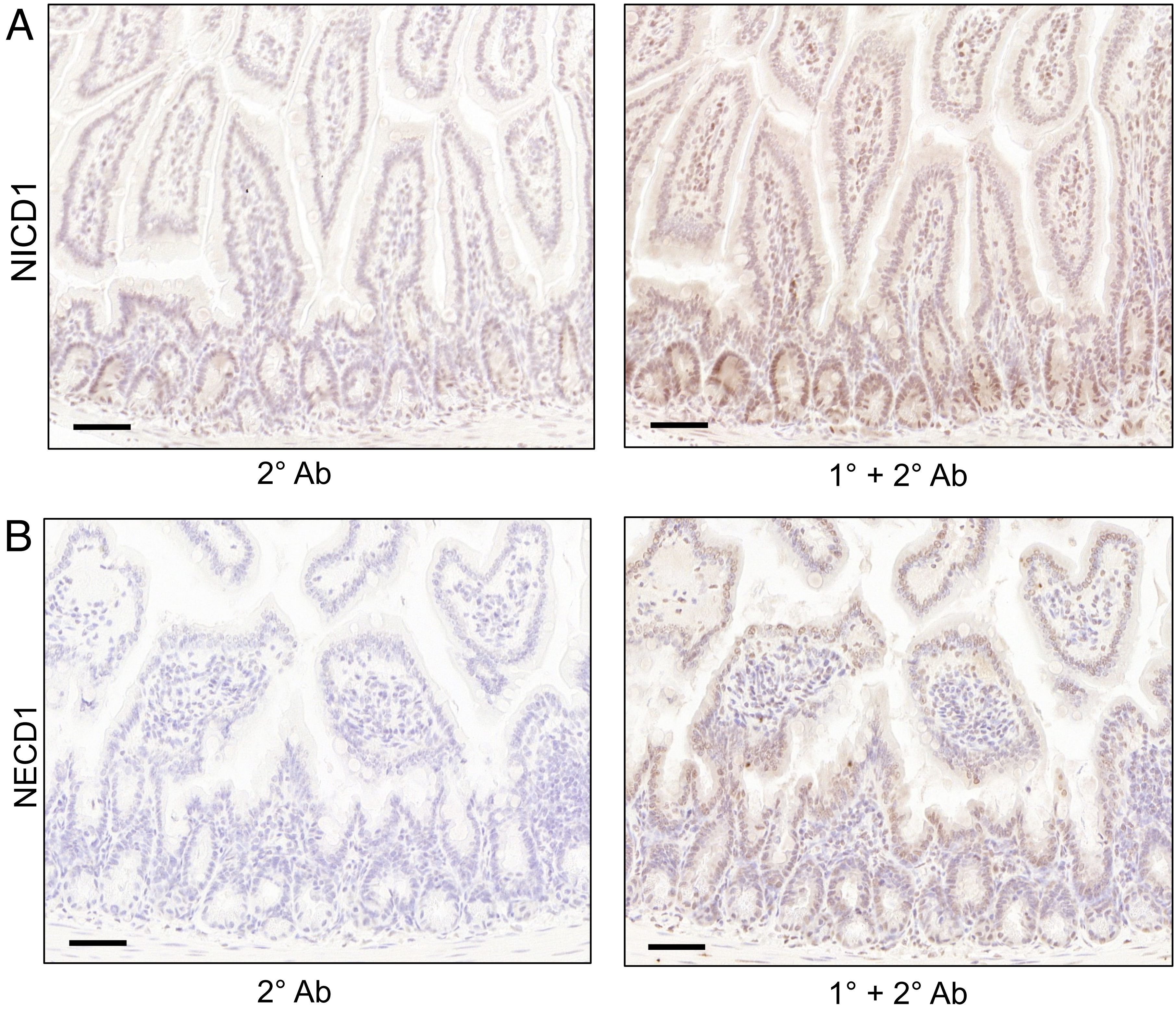


**Supplementary Figure 7.** Controls for western blotting analysis. (*A* and *B*) Small intestine tissues from Control mice showing signals with and without primary antibodies for NICD1 and NECD1 by IHC.

### Supplementary Figure 8

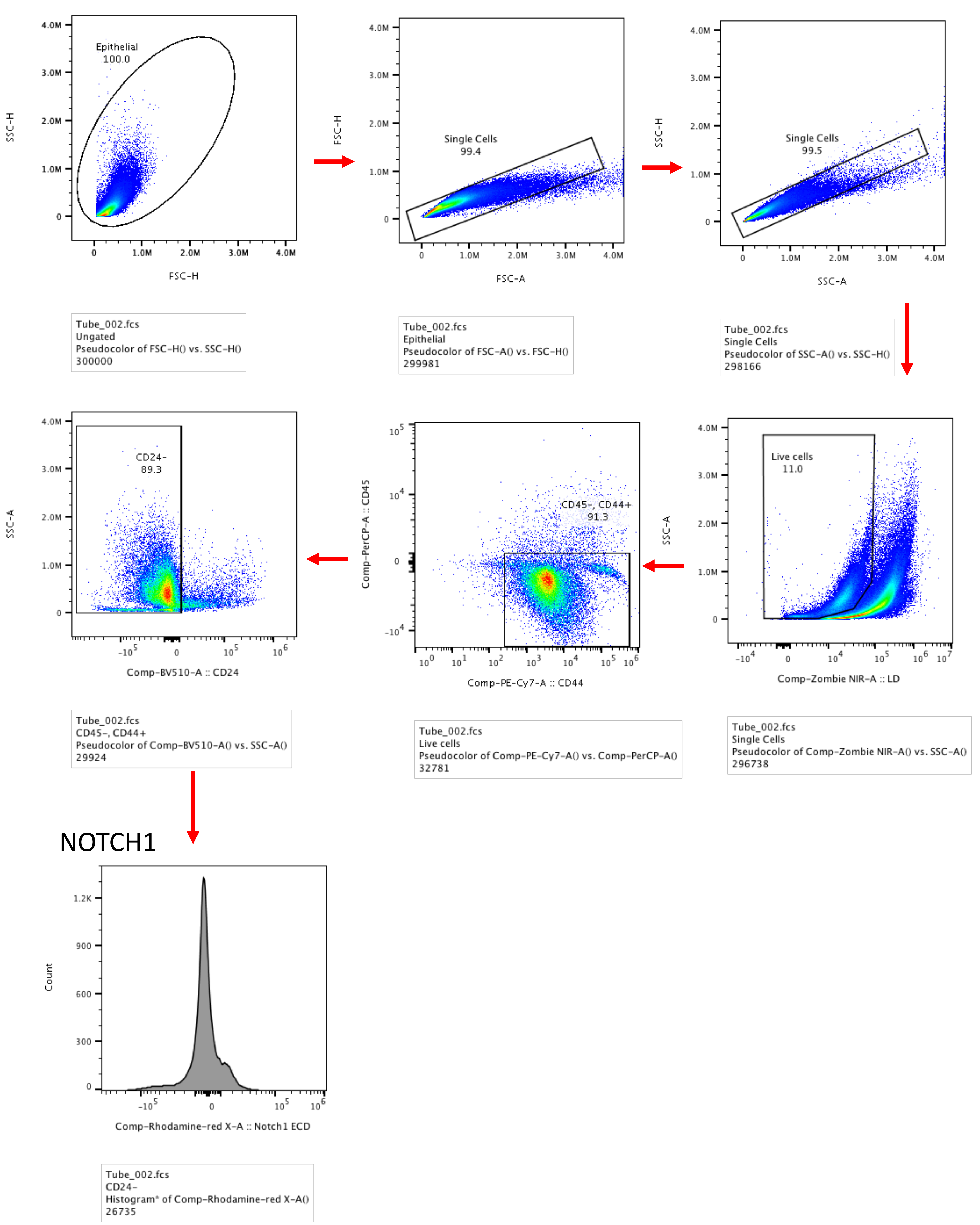

**Supplementary Figure 8.** Control mouse showing gating strategy to analyse binding of anti-NOTCH1 ECD or different soluble Notch ligands in ISC of experimental mice. Single live cells were gated to select $CD45^{-}CD44^{+}CD24^{-}$ cell population. $CD45^{-}CD44^{+}CD24^{-}$ cells were used to calculate Mean fluorescence index (MFI) for anti-NOTCH1 binding or only 2° Ab sample. Similar gating strategy was used to determine DLL1-Fc or DLL4-Fc binding to ISCs.
